# A High-Throughput Data-Independent Acquisition Workflow for Deep Characterisation of the *sn*-Isomer Lipidome

**DOI:** 10.1101/2023.11.04.565044

**Authors:** Jesse A. Michael, Reuben S. E. Young, Rachelle Balez, Lachlan J. Jekimovs, David. L. Marshall, Berwyck L. J. Poad, Todd W. Mitchell, Stephen J. Blanksby, Christer S. Ejsing, Shane R. Ellis

## Abstract

We report a workflow based on ozone-induced dissociation for untargeted characterization of hundreds of *sn-*resolved glycerophospholipid isomers from biological extracts in under 20 minutes, coupled with an automated data analysis pipeline. It provides an order of magnitude increase in the number of *sn-*isomer pairs identified compared to previous reports, reveals that *sn*-isomer populations are tightly regulated and significantly different between cell lines, and enables identification of rare lipids containing ultra-long chain monounsaturated acyl chains.

## Main

Glycerophospholipids (GPL) are one of the most ubiquitous lipid classes in biology and the most abundant in vertebrata. GPLs play many important functions within cellular membranes, including regulating membrane mechanics, signalling and protein function^1-3^, whilst changes in lipid metabolism and the lipidome are a hallmark of many diseases, including cancers^4, 5^. The biological function and metabolic origin of lipids depends on their precise chemical structure, including headgroup (defining the GPL class), the length and degree of unsaturation of each acyl chain, and the positions of each chain on the glycerol backbone (defined as the stereospecific numbering [*sn*]-position)^6, 7^.

Conventional mass spectrometry-based lipidomics workflow identify GPLs at the molecular lipid species (MLS)-level (*e*.*g*., phosphatidylcholine (PC) 16:0-20:4), but do not resolve deeper levels of structural detail such as the *sn*-position of fatty acyls (**Supplementary Figure 1**) or position(s) of double bonds without sophisticated method validation and calibration^8^. This underestimates sample complexity and poses a limitation in the biological interpretation of lipidomics data^9^. For example the relevance of *sn*-specificity to lipid metabolism is highlighted by the action of phospholipase A2 (PLA2) on membrane GPLs, resulting in the release of polyunsaturated fatty acids (PUFAs) such as arachidonic acid at the *sn*-2 position, is a well-known initiating step in inflammatory processes^10^.

Membrane lipids maintain a great diversity of acyl chain characteristics, ranging from short to long and saturated through to polyunsaturated. While historical investigations found GPLs to exhibit a preference for PUFA substitution at the *sn-*2 position (so-called canonical isomers or canonomers), advances in technologies have revealed hitherto apocryphal PUFA substitution at the *sn*-1 position (so-called apocromers)^11, 12^. These findings suggest the presence of both *sn*-isomers at varying relative abundance is expected across mammalian biology. Disruption to the spatial distribution of *sn*-isomers has been observed in cancerous brain tissue^11^, as well as individual neuronal cells^13^, suggesting different biological functions for *sn*-isomers. The ultimate purpose of such *sn*-isomer diversity is not well-understood but, in part, arises through activity of the Lands cycle, where GPLs are continuously remodelled by PLA1/2 and acyl transferase enzymes^14^. Thus, techniques capable of characterising the *sn*-isomers of lipids are needed to enhance our understanding of lipid metabolism and functions in a variety of diseases.

Traditionally the study of *sn*-isomers required phospholipase A2 assays performed on isolated lipid fractions^15 16^. Moreover, broad separation of *sn*-isomers by reversed-phase chromatography is challenging and requires long run times and extensive pallet of synthetic standards to assign peaks to individual isomers^12^. To address this challenge tandem mass spectrometry methods that resolve GPL isomers have recently been developed for deeper characterisation of the lipidome^17, 18^. While studies assigning double bond location to hundreds of distinct GPL species have been demonstrated^19^, coverage of the *sn*-isomer lipidome via techniques that provide isomer-specific fragment ions has thus far not been as comprehensive, with the most detailed studies reporting up to ∼50 *sn*-resolved species^19-21^.

Sequential collision-induced dissociation and ozone-induced dissociation (CID/OzID) is an MS^3^ technique that can resolve lipid *sn-*isomers^22^. CID of alkali-adducted GPLs leads to a neutral loss of the phosphate-containing headgroup and importantly the formation of a new carbon-carbon double bond attached to the *sn*-2 position acyl chain. Ozonolysis of this double bond subsequently produces fragments diagnostic of the FAs in the *sn*-1 and *sn*-2 positions^22, 23^ (cf. **Supplementary Figure 2**). To-date, this technique has been applied to only a small number of targeted precursors^22, 23^. Here, we report the extension of CID/OzID in an untargeted, data-independent analysis (DIA) shotgun lipidomics workflow for the deep characterisation of PC and phosphatidylethanolamine (PE) species in complex biological samples. The approach combines the specificity of 1 Da wide selection windows with MS^3^-based CID/OzID performed with the high-resolution (500,000 at *m/z* 200) and mass accuracy (<3 ppm) of a Tribrid Orbitrap mass spectrometer. The method is underpinned by an automated data analysis pipeline, that uses an updated version of ALEX^123^ for high-confidence identification of lipid species^24^ using a curated CID/OzID fragmentation library and a set of strict fragmentation criteria.

The utility of our method is demonstrated by analysis of four mammalian cell lines (HEK293T, SH-SY5Y, RAW264.7 and fibroblasts) cultured under identical conditions. Lipids were extracted^25^ and analysed in positive ion mode using nanoelectrospray ionisation in the presence of sodium formate (**Figure 1a**) to preferentially yield the required sodium adducts (**Supplementary Figure 3**). Central to the method is a DIA approach by which lipid precursor ions in the mass range of *m/z* 574-1036 are sequentially selected in ±0.5 wide windows and subject to CID/OzID-MS^3^ analysis (**Figure 1b**). Individual precursors are first fragmented using CID, with fragment ions corresponding to the neutral loss of *m/z* 183.0661 Da (from PC lipids) or 141.0191 Da (from PE lipids) being re-isolated in the linear ion trap and exposed to ozone for 400 msec. Sequentially stepping through the indicated *m/z* range ensures all PE and PC species are subjected to CID/OzID analysis, including low abundance species that would not be interrogated by DDA approaches. The data is processed through the ALEX^123^ pipeline, with peaks being matched to an extensive *in-silico* spectral library featuring CID/OzID-MS^3^ fragments for all possible *sn*-positional GPL isomers with FA chains that range from 12 to 36 carbons (**Figure 1c**). This is followed by automated identification procedure where fragment assignments are subjected to a stringent ±2.5 ppm mass accuracy threshold (see methods). The high-confidence identification approach is validated through analysis of a wide assortment of synthetic lipid standards to elucidate how different acyl chains alter which specific CID/OzID fragments are preferentially formed (**Supplementary Figure 4)**.

**Figure 1.**
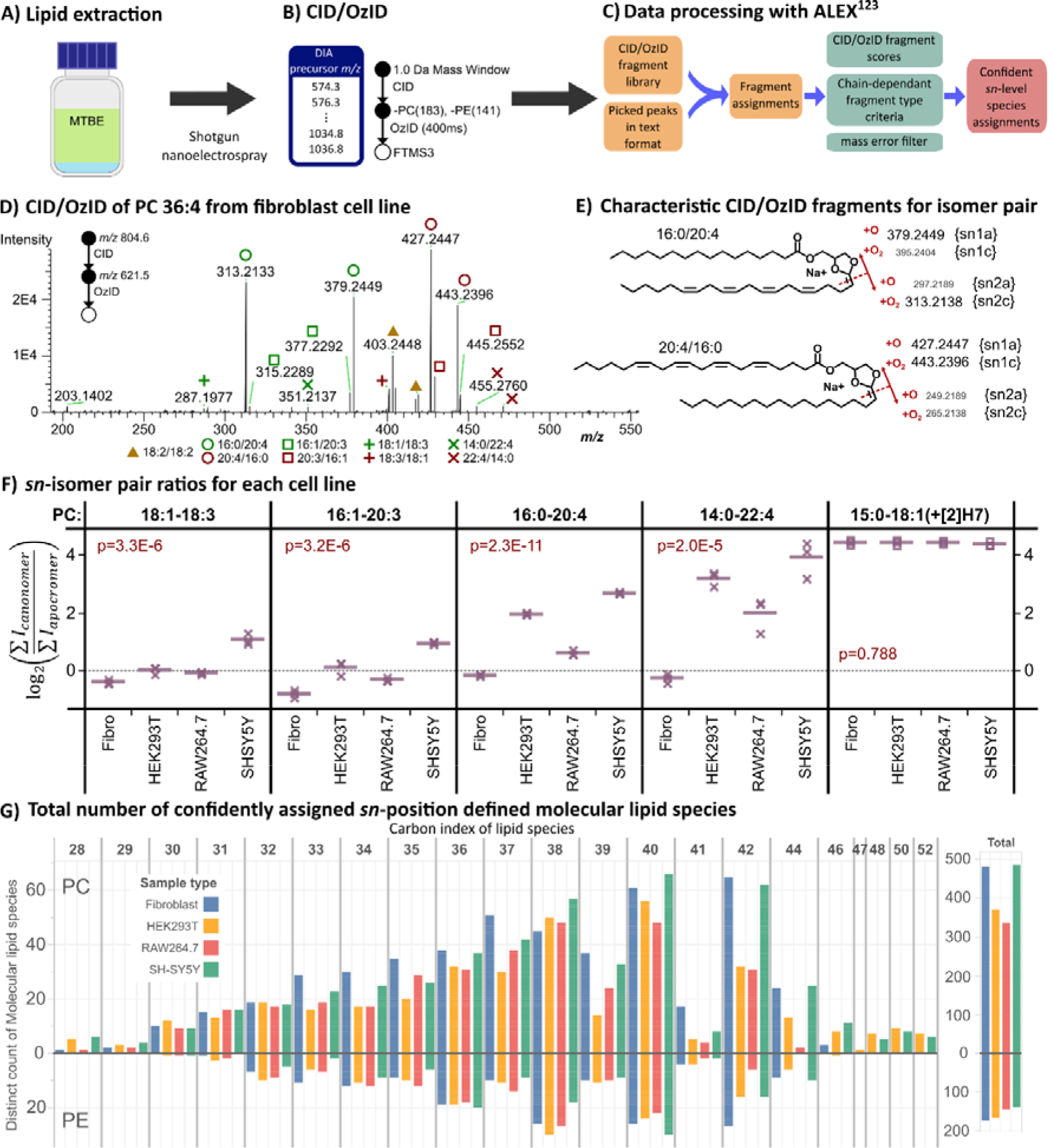
**a, b, c**, Summary of the automated DIA-CID/OzID workflow. **d**, CID/OzID spectrum acquired from fibroblasts determining the *sn*-isomer pairs for the PC 36:4 reveals four isomer pairs. **e**, CID/OzID fragmentation of PC 16:0/20:4 and PC 20:4/16:0 with fragments indicated. **f**, Log_2_ ratios of fragment ion intensities for four *sn*-isomer pairs detected from PC 36:4 for each of the cell lines, along with the internal PC standard (far right column). Mean fragment ion intensity ratios are represented by the line and individual replicates marked with crosses. p-values for a 1-way ANOVA test for differences in isomer ratio between cell types are given. **g**, Total number of distinct PE and PC molecules that were confidently annotated in each cell line, broken down by carbon index of lipid species.

From fibroblasts, CID/OzID of PC 36:4 results in confident annotation of 9 distinct lipid molecules, including four *sn*-isomer pairs (**Figure 1d**). The product ions expected for the dominant *sn-*isomer pair, PC 16:0/20:4 and 20:4/16:0, are shown in **Figure 1e** (green circles and squares), which illustrates that these lipids are each represented by two main isomer-specific fragments. The ratio of peaks relating to the canonomer (*i*.*e*., the expected isomer with the longer and more unsaturated chain at the *sn*-2 position; PC 16:0/20:4) to apocromer (*i*.*e*., the inverse conformation; PC 20:4/16:0) was found to be reproducible across replicates but significantly different between the cell types (**Figure 1f, Supplementary Figure 5**). For comparison, the synthetic internal standard (PC 15:0-18:1(^2^H_7_), composed of PC 15:0/18:1(^2^H_7_) and PC 18:1(^2^H_7_)/15:0, showed no significant difference in isomer ratio, demonstrating the specificity of the approach to biological changes in isomer populations.

The sensitivity and coverage of the DIA-CID/OzID workflow is exemplified in **Figure 1g** whereby up to 508 *sn*-resolved PC species and 144 *sn*-resolved PE species were detected in each cell line. The lower number of PE annotations is attributed to their lower absolute abundance. In total 966 unique diacyl PC and PE lipids were observed with 296 being common amongst all four cell lines (**Supplementary Data 1**). Despite being of low abundance, 34% of PC and 25% of PE annotations contained odd-numbered FA chains. In addition, 136 ether-linked PEs and PCs were identified (**Supplementary Data 2**). We validated the identified FAs by fatty acid profiling using LC-MS analysis of hydrolysed and AMPP-derivatised fatty acyl chains^26^ from SH-SY5Y cells, verifying ≥97% of all FA assignments (**Supplementary Data 3**).

The potential of our method for *de novo* discovery is exemplified by the detection of rare PC molecules containing ultra-long-chain (ULC) FAs consisting of 26-36 carbons (**Figure 3a-b**). While several 26 carbon-containing PUFA species were observed only in fibroblasts, a series of ULC-monounsaturated fatty acyls (ULC-MUFAs) up to 36 carbons were only observed in SH-SY5Y and HEK293T cells (**Figure 2, Supplementary Figure 6)**. A representative CID/OzID spectrum for PC 50:2 is shown in **Figure 2a** with annotated fragments for PC 16:0/34:2, PC 18:1/32:1, PC 32:1/18:1 and PC 34:1/16:1. **Figure 2b** shows the FA combinations and relative signal intensities for all ULC-containing species, revealing the strong preference for esterification of the ULC-MUFAs to the *sn*-1 position. While ULC-PUFAs in the *sn*-1 position of PCs have been well-described in mammalian tissues^27, 28^, we find few reports of ULC-MUFAs in GPLs^29, 30^. Several of these molecules, including the relatively abundant PC 32:1/18:1 and PC 34:1/16:1, to the best of our knowledge, are novel lipids. The presence of the ULC-MUFA chains were also validated by LC-MS analysis of AMPP-derivatised hydrolysed fatty acyls (**Supplementary Figure 7)**, which also identified their double bond position to be majority *n*-9 (**Supplementary Figures 8-10**). We hypothesise these FA chains parallel the described synthesis of ULC-PUFAs and arise from elongation of oleic acid (18:1*n-*9) by ELOVL4^6, 27, 29^.

**Figure 2.**
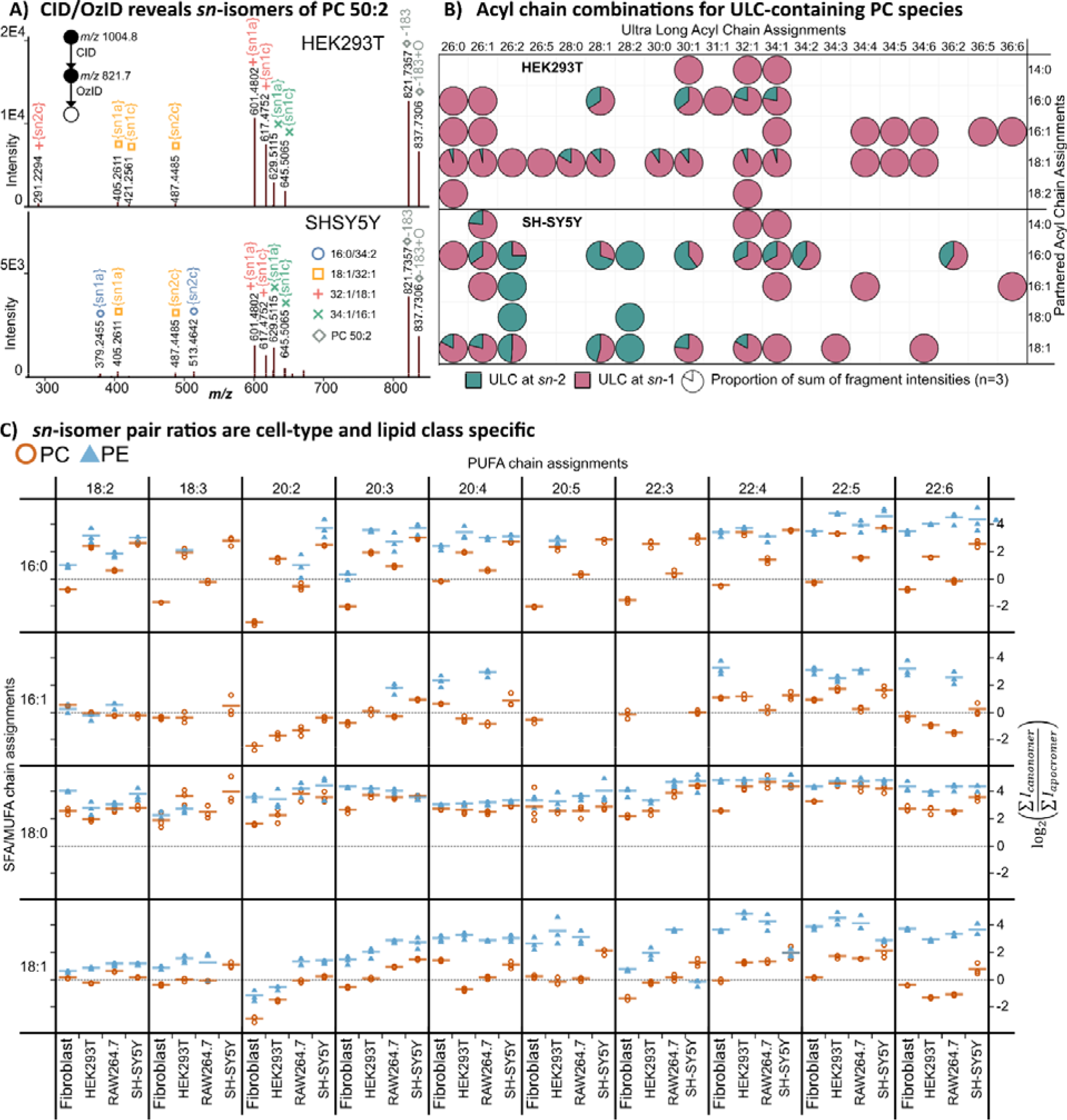
CID/OzID identifies novel lipid species containing ultra-long chain (ULC) monounsaturated fatty acyls (ULC-MUFAs) present with a high specificity towards the *sn*-1 position of PCs, as well as revealing how *sn*-isomer populations vary across cell lines. **a**, CID/OzID spectra of PC 50:2 revealing the presence of 32:1, 34:2 and 34:1 chains in HEK293T and SH-SY5Y cells present predominantly at the *sn*-1 position. **b**, Presence of ULC-containing PC lipid isomers identified across HEK293T and SH-SY5Y cell types, where pie fractions are proportional sum of product ion intensities from three replicates for each isomer. **c**, PC and PE *sn*-isomer fragment ion ratios for selected lipids containing different combinations of either saturated fatty acyls (SFA) or monounsaturated fatty acyls (MUFAs) and a polyunsaturated acyl chain (PUFA) across fibroblast, HEK293T, RAW264.7 and SH-SY5Y cell types. Each mark represents the *sn*-isomer ratio calculated for a single replicate, with bars indicating the mean of 3 replicates. Canonomer refers to the expected molecule with the more unsaturated chain at the *sn*-2 position and apocromer refers to the unconventional molecule containing the more unsaturated chain at the *sn*-1 position. Note that the isomer pair ratios calculated by fragment intensities do not necessarily reflect mole fractions of isomeric populations.

Importantly, our method permits extensive comparison of *sn*-isomer populations across biological systems. **Figure 2c** displays the isomer ratios from all cell lines for selected lipids with common SFA-PUFA and MUFA-PUFA chain combinations. Although the proportion of saturated and monounsaturated lipids have been shown to be reflected by CID/OzID analysis with high fidelity^22, 31^, extensive unsaturation imparts differing ozonolysis reaction rates to different isomers meaning fragment ion abundances do not necessarily reflect mole fractions of isomers. Accordingly, *sn-*isomer ratios across the four cell lines were compared which provides insight into relative changes of isomer populations for a given *sn*-isomer pair. Notable differences in the abundance of *sn-*isomers could be observed within both lipid class and specific FA chain combinations. For example, PC species frequently had a higher apocromer population than the similar PE species with equivalent FA chains. Concurrently, lipids from fibroblasts were observed to have higher apocromer fractions compared to other cell lines for species containing FA 16:0, however, this was not consistent for species containing 16:1, 18:0 or 18:1 chains. These results suggest that cells have fine control over *sn-*isomer metabolism.

In summary, our method overcomes a major limitation of current lipidomic workflows and permits deep and automated characterisation of the PC and PE lipidome with *sn-*isomer resolution in ∼20 minutes/sample. It provides new molecular insight into the cell-specific regulation of lipid isomer populations in biological samples and enable the discovery of novel lipid molecules, specifically PC and species with ULC-MUFA chain at the *sn*-1 position. Moreover, the approach has increased the depth of *sn-*isomer coverage by an order of magnitude compared to previous methods and our automated data processing pipeline enables interactive data visualizations to inspect fragment assignments, perform quality control and compare samples. The method could be improved in the future by developing derivatisation methods to enable positive mode analysis of further GPL classes. While the workflow here is validated for widely used Orbitrap mass spectrometers, the ALEX^123^ data processing pipeline can be amended to handle data files from a variety of instrument platforms that allows MS^3^ or pseudo MS^3^ and OzID such as the orthogonal time-of-flight-based platforms.

## Acknowledgements

J.A.M. acknowledges support from the Australian Government Research Training Program Scholarship. S.R.E. acknowledges support from the Australian Research Council Future Fellowship Scheme (FT190100082) and, along with R.S.E.Y., the Human Frontiers Science Program Organization (RGP0002/2022). The coupling of the Nanomate to the orbitrap Fusion was enabled by a University of Wollongong Small Project Grant. S.J.B. and T.W.M. acknowledge funding through the Australian Research Council Discovery Program (DP190101486). C.S.E. acknowledges support from the VILLUM Foundation (VKR023439), the VILLUM Center for Bioanalytical Sciences (VKR023179) and European Molecular Biology Laboratory (EMBL). We are grateful to the technical support provided by the University of Wollongong Mass Spectrometry Facility.

## Methods Cell culture

Four cell lines, HEK293T: human embryonic kidney; SH-SY5Y: human neuroblastoma; RAW264.7: murine macrophage, and fibroblasts, a non-immortalised human dermal fibroblast line, were cultured under identical conditions. All cell lines were maintained in Dulbecco’s modified Eagle medium (DMEM) High Glucose, supplemented with 10% (v/v) heat-inactivated fetal calf serum (Interpath FBS-F, Australia) at 37 °C in a humidified incubator with 5% CO_2_. Medium was changed every 3-5 days. To harvest, cells were washed with phosphate buffered saline (PBS) and incubated with 0.5% Trypsin/EDTA (Life Technologies, USA) for 3-5 min. Trypsin was inactivated with the addition of DMEM High Glucose + 10% FCS and cells were aspirated and centrifuged at 300 × g for 5 min. Cells were re-suspended in 1 mL PBS for counting and then washed with 10 mL PBS and centrifuged at 300 × g for 5 min. The PBS was aspirated and the cell pellet re-suspended in 100-300 *μ*L LCMS grade methanol (VWR International, Australia) with 0.1% butylated hydroxytoluene (BHT) (Sigma-Aldrich, Australia) and stored at -80 °C until use.

### Cell culture lipid extractions using modified MTBE method

For each cell sample, 1-2 x 10^6^ cells in methanol with 0.1% BHT were combined with EquiSplash internal standard mixture (Avanti Polar Lipids, Alabaster, AL, USA) (2 *μ*L per 1 x 10^6^ cells) and made up to a total volume of 400 *μ*L using methanol, which was homogenized using a Fastprep-24 (MP Biomedicals) with two rounds of 6 m/s for 30 s with resting in ice for 2 minutes in between. 240 *μ*L of homogenate was aliquoted to a 1.5 mL glass-vial containing 800 *μ*L of MTBE. Vials were then vortexed for 1 hour at room temperature. Subsequently 200 *μ*L of aqueous 100 mM ammonium formate was added, the sample vortexed for 20 s and then centrifuged for 5 minutes at 750g. The top (MTBE) phase was collected and stored at -20°C. For mass spectrometry analysis, MTBE phases were aliquoted into 1.5 mL Eppendorf tubes and vacuum evaporated using a SpeedVac SC10 (Savant Instruments Inc.) before reconstitution in isopropyl alcohol/methanol (2:1 vol./vol.) containing 120 *μ*M sodium formate and added to a 96 well-plate which was foil heat-sealed and centrifuged.

### DIA-CID/OzID mass spectrometry analysis

Shotgun lipidomics using DIA combined with CID/OzID was performed in positive-ion polarity using an Orbitrap Fusion Tribrid mass spectrometer (Thermo Scientific, San Jose, CA, USA) coupled to a TriVersa NanoMate (Advion Biosciences, Ithaca, NY, USA). OzID was performed using a high-concentration HC-30 ozone generator (Ozone Solutions) and doping the ozone output into the helium supply of the linear ion trap (LIT) through a PEEK restriction, similar to that described previously^11, 22^. The ozone generator was supplied with 0.2 L/min of oxygen and output 130-140 g/m^3^ ozone concentration (ca. 10% ozone in oxygen), as recorded by an in-line ozone monitor. Parameters for the NanoMate were 1.15 kV spray voltage, 1.15 psi pressure with tip pre-wetting, mandrel foil piercing and the sample stage held at 10°C.

A DIA acquisition, designed in the Thermo Xcalibur method editor, was used to collect MS^1^, MS^2^ and MS^3^ (CID/OzID) scans over a 35 min duration with the MS^3^ *sn-*isomer data for PC and PE species taking ∼20 min (the only data used here for *sn*-isomer characterisation). MS^1^ spectra were collected for *ca*. 8 minutes with an *m/z* range of 350-1600 at a target mass resolution of 500,000 at *m/z* 200 (FWHM) and an automatic gain control (AGC) at 50% (manufacturer units) and a 50 ms maximum injection time. MS^2^ activation used resonant CID (MS^2^) with a nominal collision energy of 40%, 5 ms activation time, activation q value of 0.25 and ±0.5 Da isolation window. MS^2^ spectra were acquired using a target mass resolution of 120,000 at *m/z* 200 with a maximum injection time of 246 ms and an AGC target of 300%. MS^3^ (CID/OzID) scans were recorded using a target list containing all even *m/z* values between 574.3-1036.8 for MS^2^ activation and an *m/*z value 183.0661 or 141.0191 Da below the precursor ion used for MS^3^ activation (for PC and PE lipid classes, respectively). MS^3^ activation used an isolation window of ±0.6 Da, an AGC target value of 300% (manufacturer units), a collision energy of 0%, an (ozonolysis) activation time of 400 ms, a maximum injection time of 1012 ms, and Orbitrap detection at a target mass resolution of 500,000 at *m/z* 200. One scan for each value in the target list was acquired per sample.

### Preparation of fatty acids derivatives

Fatty acid analysis was conducted on the hydrolysed cell lipid extracts using the N-(4-Aminomethylphenyl) Pyridinium (AMPP) derivatisation method first described by Bollinger *et al*^*32*^. and later adapted by Young *et al*^*33*^ and Menzel *et al*^*26*^. Briefly, 50 *μ*L of lipid extract was added to 4 mL glass vials and dried under nitrogen gas flow. Fatty acids were hydrolysed from intact lipids using 200 *μ*L tetrabutylammonium hydroxide (40% w/v water) and 200 *μ*L methanol with heating (75 °C) for 2 h. Samples were allowed to cool to room temperature before adding 1.5 mL of cold n-pentane, 1.5 mL of water and 300 *μ*L of aqueous 5 M hydrochloric acid before vortexing for 30 sec and supernatant collection. 150 *μ*L of acetonitrile/dimethylformamide (4:1 v/v) was added to the collected supernatants before vortexing and removal of the volatile organic solvents under a stream of nitrogen gas. An additional 150 *μ*L of acetonitrile/dimethylformamide(4:1 v/v) was added to the residual solvent droplet post nitrogen gas drying and the vial was vortexed for 30 sec. 5 *μ*L of freshly prepared EDC solution (20 mg 1-ethyl-3-(3-dimethylaminopropyl)carbodiimide hydrochloric acid in 100 *μ*L of water) was added before adding 40 *μ*L of HOBt solution (30 mM 1-hydroxybenzotriazole in acetonitrile:dimethylformamide(4:1 v/v)) and vortexed for 30 sec. 40 *μ*L of AMPP solution (20 mM in acetonitrile) was added to the mixture and the vial contents was heated at 65 °C for 30 min. After allowing to cool to room temperature, 1 mL of MeOH was added and volatiles were removed under nitrogen gas flow. The dry residue was reconstituted in 1 mL of methanol for LC-OzID-MS analysis.

### LC-OzID-MS analysis of AMPP derivatised fatty acids for fatty acyl validation

For the confirmation of fatty acyl species observed within the described CID-OzID analysis, LC-OzID-MS analysis of hydrolysed fatty acids was undertaken using the ozone-induced dissociation fatty acid discovery (OzFAD) workflow^26^, with minor adaptations. Briefly, analysis was performed using a Waters Acquity (UPLC i-Class; CSH, C18 reversed phase column, 2.1□×□100□mm, particle size 1.7□*μ*m) liquid chromatography system coupled with a Waters SYNAPT G2-Si (Z-Spray, T-Wave Ion Mobility; TOF) mass spectrometer that had been modified to perform OzID experiments^34^. Ozone was generated from UHP oxygen by an ozone generator (Ozone solutions, TG-40; generating 200–250□gm^−3^ of ozone in oxygen) and introduced to the ion mobility (IMS) cell at a flow of 90□mL□min^−1^ in place of the default nitrogen gas. Using the derivatised cell fatty acids samples described (*vide infra*), 5 *μ*L of sample was injected. Chromatographic separation of the sample analytes was performed using an oven temperature of 60 °C, a flow rate of 0.4□mL□min^−1^, and a two-mobile phase system (mobile phase A: water containing 0.1% formic acid; mobile phase B: acetonitrile containing 0.1% formic acid). The mobile phase mixing gradient was as follows (in percentages of mobile phase B): 0.00 - 0.50 min, isocratic at 20%; 0.51 - 16.00 min, ramp from 20% to 90%; 16.01 - 21.00 min, ramp from 90% to 100%; 21.01 - 24.00 min, isocratic at 100%; 24.01 - 24.02 min, ramp from 100% to 20%; 24.02-24.50 min, isocratic at 20%. The mass spectrometer was operated in positive polarity, sensitivity and IMS-modes. For high energy CID experiments, the quadrupole was operated in RF-only mode (MS method: MSe Continuum, 30-100 eV collision energy applied in the transfer region of the instrument, post OzID). Ozonolysis was performed using the instruments IMS mode, which allows for ∼15 ms of analyte exposure to ozone. The resulting monoisotopic ions (precursors and products) were then mass analysed by time of flight (nominal resolution *m/Δm* ∼ 15,000). Sample data was manually investigated using the Waters MassLynx software (version 4.2), with positive assignment of AMPP FA species requiring (i) identification of the AMPP head group fragment (within the high energy experiment), (ii) the precursor ion Δ*m/z* being within ±0.005 of the theoretical value, and (iii) a minimum signal-to-noise ratio of three. Double bond assignments additionally required the retention time of the identified AMPP FA precursor ion and the aldehyde and Criegee product ions to be temporally aligned within the chromatogram. Note that due to the presence of PC 15:0-18:1(^2^H_7_) and PE 15:0-18:1(^2^H_7_) internal standards endogenous lipids containing 15:0 acyl chains were excluded from the validation procedure.

### Lipid identification from DIA-CID/OzID data using ALEX^123^

Lipid identification was carried out using ALEX^123^ software, which has been described and is publicly-available^24^. For identification of lipid molecules detected by CID/OzID analysis, we *in-silico* generated a spectral library featuring all characteristic CID/OzID fragments that can be generated from an array of more than 100,000 *sn*-position-defined PC, PE, PC-O and PE-O species having FA chains and/or ether-linked chains with 12 to 36 carbons and 0 to 7 double bonds.

CID/OzID fragments detected by the DIA-FTMS^3^ analysis, and across all spectral datafiles (in proprietary .raw format), were matched to fragment *m/z* values in the CID/OzID library using an initial *m/z* tolerance of ±0.003 Da and an *m/z* offset of -0.0012 Da. To correct for systematic FTMS3 *m/z* calibration drifts between experiments and to determine the ppm mass error more accurately we processed the initial search results using a script (written in SAS Enterprise Guide 8.2) that automatically corrects the FTMS^3^ *m/z* calibration offset by computing the median *m/z* offset for all detected fragments across each datafile, and using this estimated calibration *m/z* offset to correct for calibration drifts between analysis and improve the mass accuracy (**Supplementary Figure S11**). Following this procedure, only CID/OzID fragments detected with a corrected mass error inside a strict ±2.5 ppm tolerance were used for further analysis.

For high-confidence identification of *sn*-position-defined molecular PC and PE species we devised a scoring algorithm that outputs a so-called CID/OzID fragment score. This scoring function is also implemented as an automatic and executable SAS Enterprise Guide script. This score function considers the number of expected fragment occurrences across all replicate analyses of a sample group. Lipids with different degrees of unsaturation and diacyl vs ether lipids, are assigned different expected numbers of fragments among the common four fragment subtypes, based on reliable patterns in fragment abundance observed from analysis of an array of standards (**Supplementary Figure 4**). For example, identification of PE 18:1/20:3 considers only the number of observations of {sn1a} and {sn2c} fragments, PE 18:3/20:1 considers only the number of observations of {sn1a} and {sn1c} fragments, while PE O-16:0/20:4 considers only the number of observations of {sn1a}, {sn2a} and {sn2c} fragments (fragment nomenclature is defined in Supplementary Figure 2). All lipids require observations of the {sn1a} fragment, detection of the CID/OzID-MS^3^ precursors (i.e. ‘-PC(183)/-PE(141)’ and the intermediate product ion ‘-PC(183)+O/-PE(141)+O’) in the same MS^3^ spectrum, as well as a CID/OzID fragment score of at least 0.8 to be considered ‘high confidence’.

We note that some minor fragments outside the dominant CID/OzID fragments described here, observed in the analysis of lipid standards, can occasionally contribute to false-positive lipid identification. For instance, analysis of PC 16:0/16:0 produces all 4 expected CID/OzID characteristic fragments, with the typical {sn1a} fragment at *m/*z 379.245 as the base peak, but with additional peaks at 377.230 and 351.251 that would be isomeric with the {sn1a} of PC 16:1/16:0 and PC 15:0/20:4, respectively. To mitigate false-position identifications, {sn1a} assignments are subject to a cut-off of 1% of the maximum {sn1a} and {sn1c} fragment in the same spectrum, or 2% and 4% of the intensity of two specific confounding fragments ({sn1a+2H} and {sn1a+CO}, respectively), which are included in the spectral library.

The SAS-based script automatically outputs two .tab files containing a summary of confident diacyl species and ether-linked species found in each sample type, as well as a detailed file containing all fragments, including those belonging to “low” and “intermediate” confidence species for quality control purposes. The detailed file can be visualised using Tableau (or other many other software platforms) with an interactive dashboard to show how fragment assignments and isomer vary across samples.

