## Supporting Information Figures for "A High-Throughput Data-Independent Acquisition Workflow for Deep Characterisation of the *sn*-Isomer Lipidome"

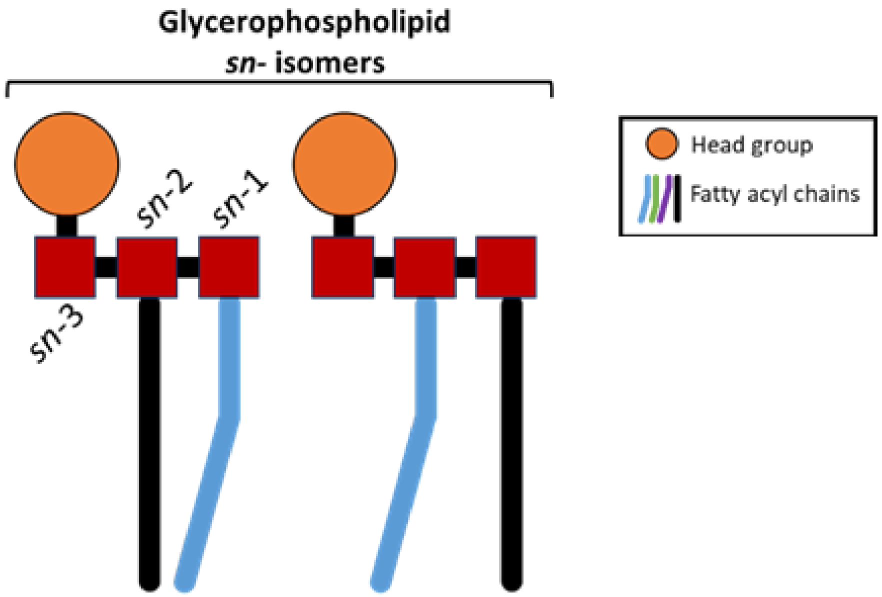


**Supplementary Figure 1.** Schematic representation of glycerophospholipid *sn*-isomers.

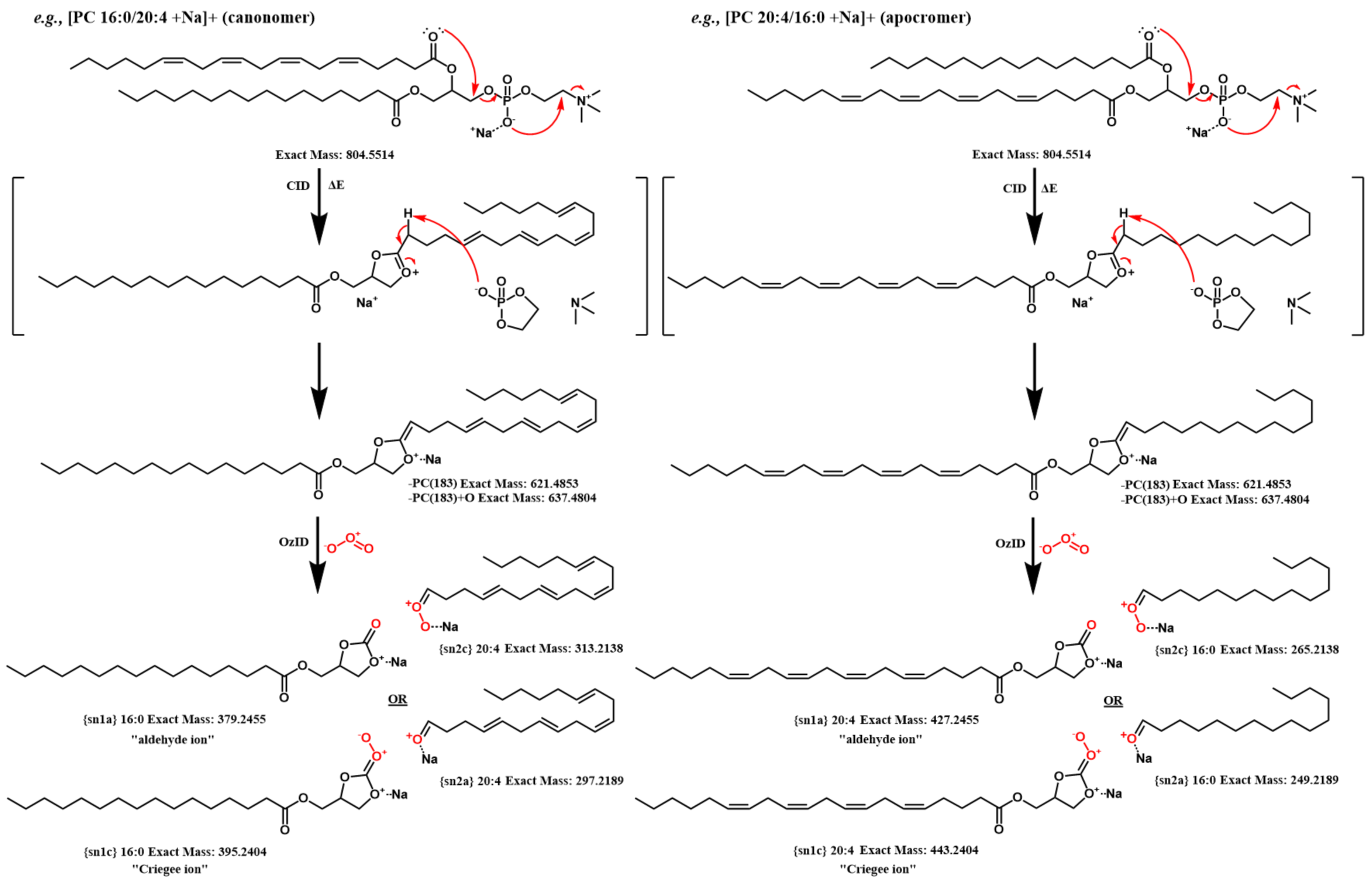


**Supplementary Figure 2.** CID/OzID mechanism and utilised fragment nomenclature for the representative species PC 16:0/20:4 (canonomer) and PC 20:4/16:0 (apocromer). Analogous process also occurs for PE lipids resulting instead in the formation of a -PE(141) ion.

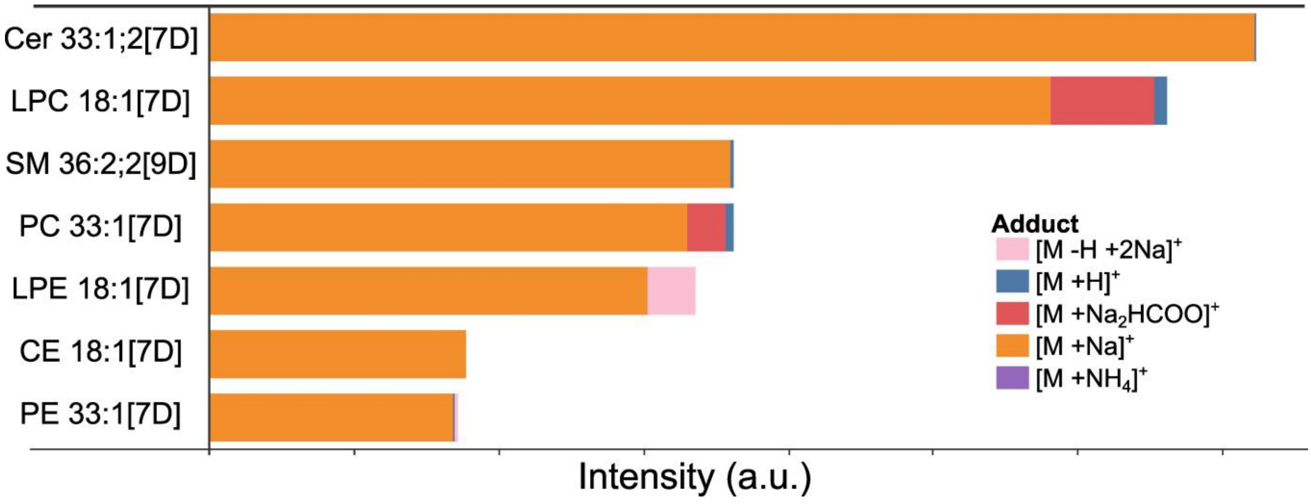


**Supplementary Figure 3.** Intensities of peaks in MS1 data corresponding to recovered equiSPLASH internal standards in lipid extracts of cell lines, demonstrating the high proportion of sodium adducts obtained. Vacuum evaporation of the extracts was found to reduce protonated adduct formation, potentially by removing residual ammonium formate present from the MTBE lipid extraction.

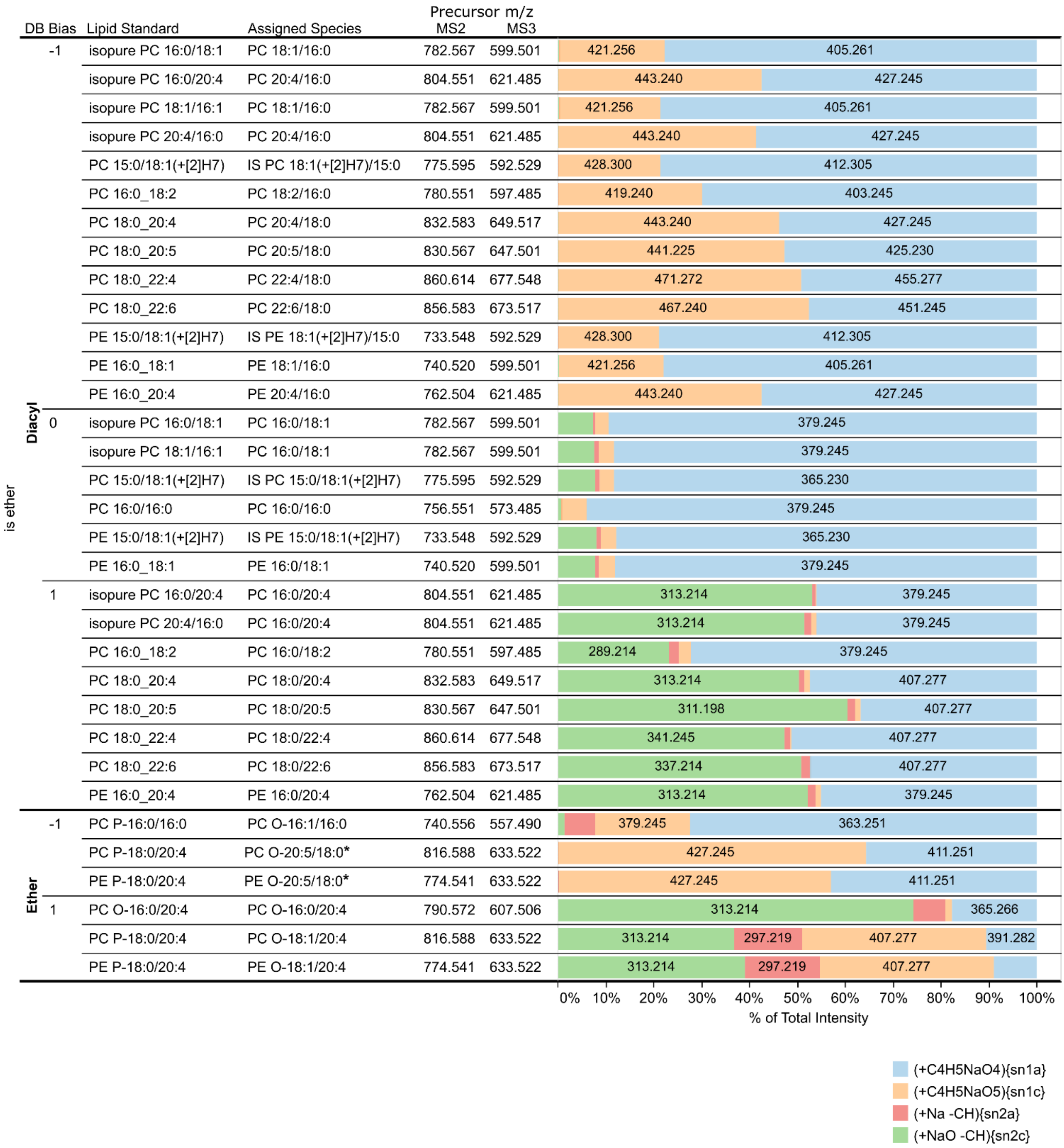


**Supplementary Figure 4.** Characteristic CID/OzID fragments observed from analysis of lipid standards presented as percentage of total fragment signal for that species. All diacyl lipid standards, including isopure standards presented fragments arising from both *sn*-isomers. Species are categorised by their "Double Bond Bias", where "-1" and "1" indicates double bonds are weighted towards the *sn*-1 or *sn*-2 chain respectively, or have a balanced distribution ("0"). These categories have notable distinctions in terms of which particular fragments are most abundant, which informs the decision-making process for de novo assignment of species in complex mixtures.

* These species are false annotations produced by ozonolysis of the plasmenyl double bond, a unique behaviour of plasmalogen species.

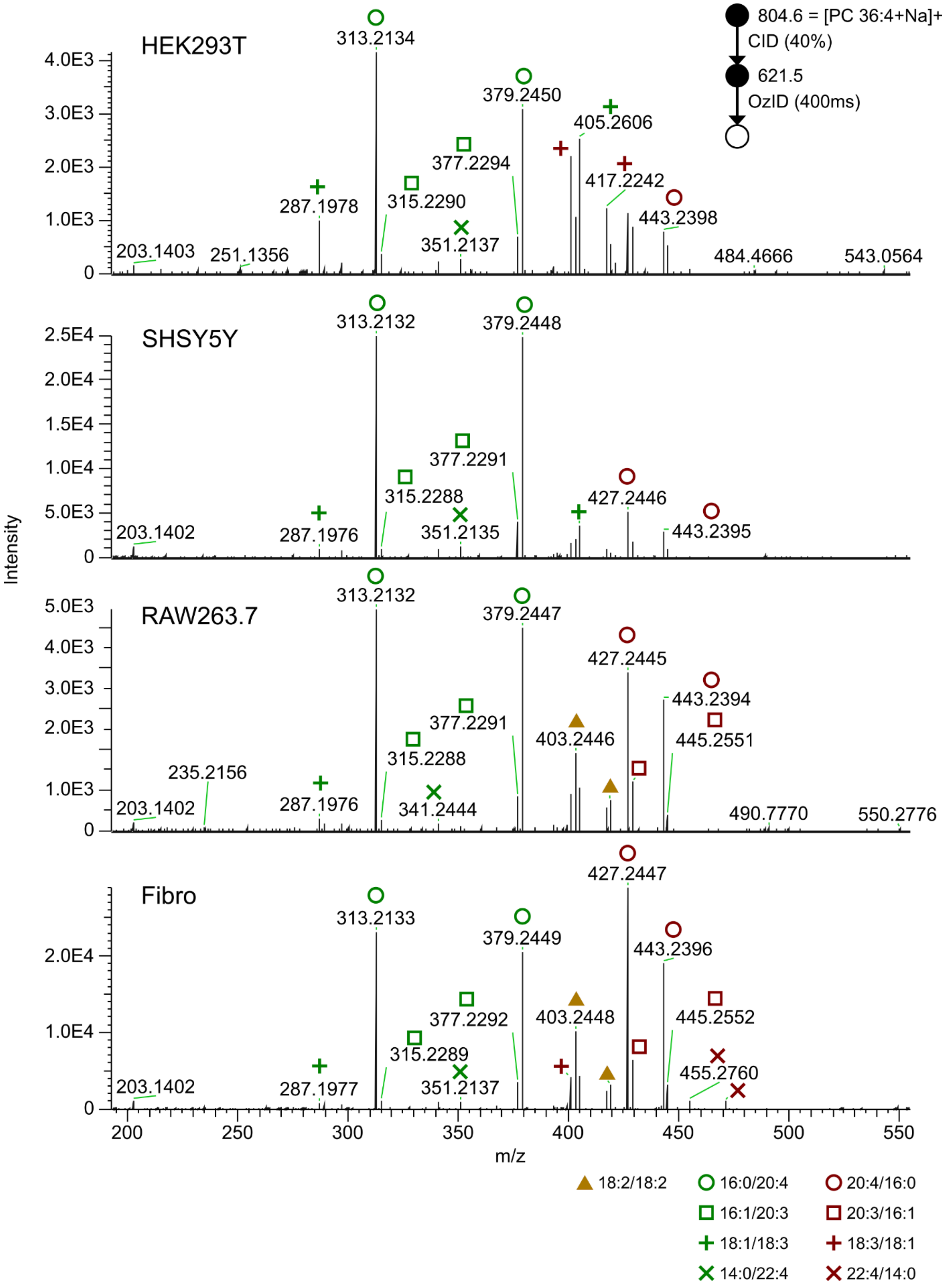


**Supplementary Figure 5.** CID/OzID spectra for PC 36:4 obtained for one replicate of each of the four cells lines. Peaks are annotated according the *sn*-resolved lipid species they correspond to. In this example four distinct isomeric pairs are observed with abundances that depend on cell type.

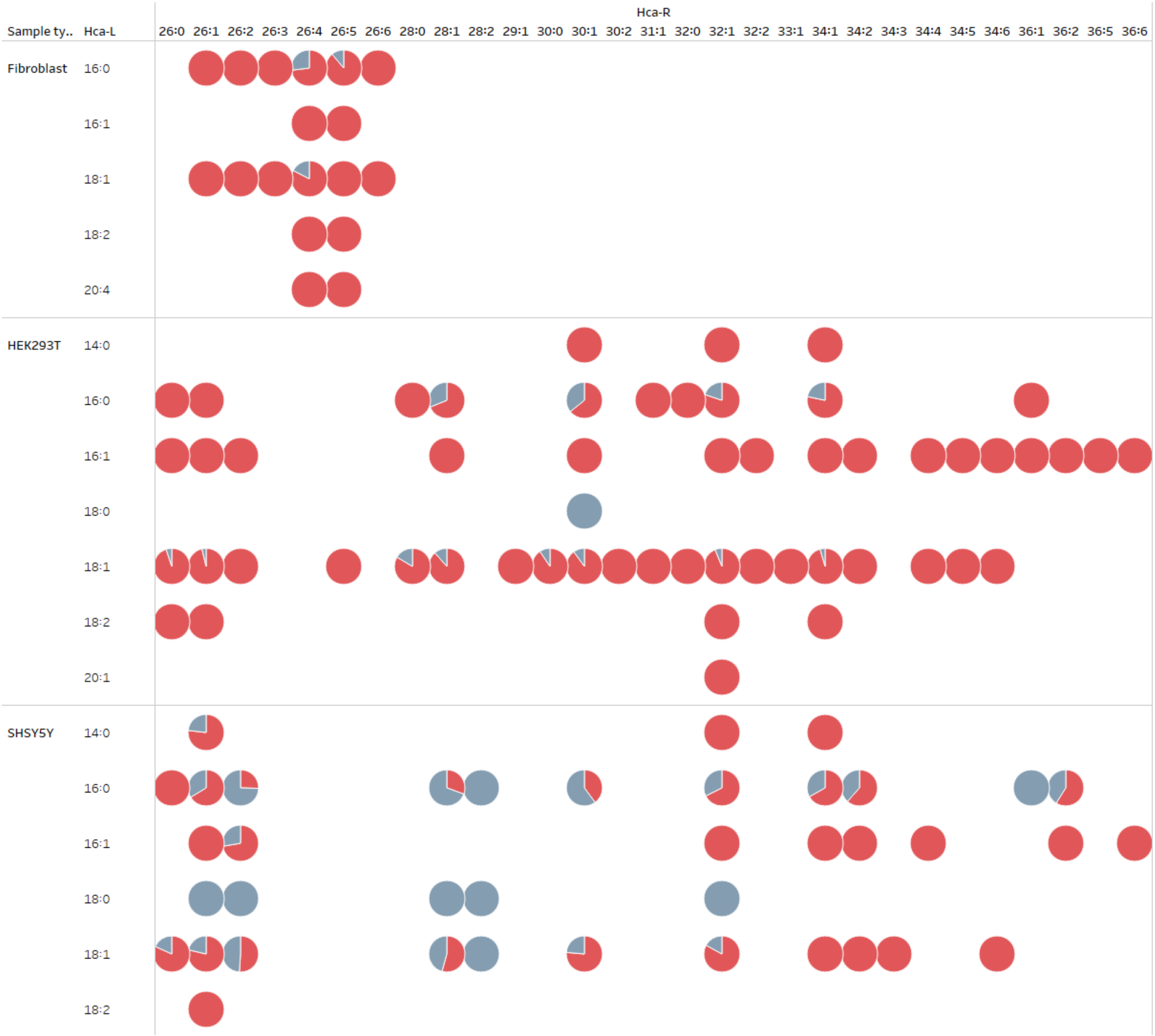


**Supplementary Figure 6.** Distributions of *sn*-isomer populations of PC species containing ultra long acyl chains (≥26 carbons) in each cell type. Each pie chart shows the relative fragment ion signal from the two *sn* isomers containing the combination of acyl chains denoted by the horizontal and vertical axes. Red and blue coloured pie segments indicate the case where the ULC is esterified to the *sn*-1 or *sn*-2 position, respectively.

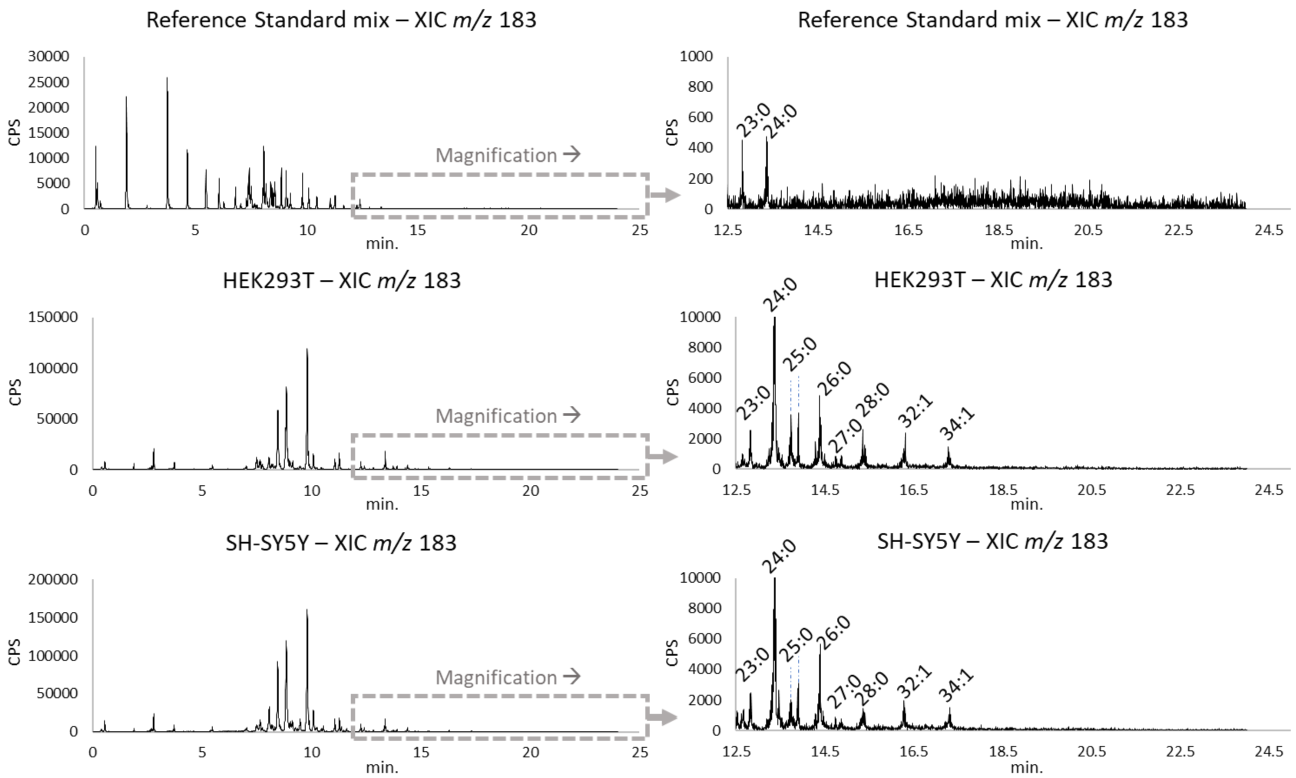


**Supplementary Figure 7**. Positive polarity LC-MS of AMPP derivatised fatty acids (AMPP-FA) from the hydrolysed lipids of (top) a synthetic reference standard mixture of 37 fatty acids, (mid) the HEK293T cell lines, and (bottom) the SH-SY5Y cell line. Using a collision energy ramp, the chromatographically separated AMPP-FA cations were fragmented using an $MS^{all}$ approach. An extracted ion chromatogram (XIC) of the fixed positive- charge headgroup fragment (*m/z* 183) was then monitored (*i.e.*, left-hand plots) and signifies the presence of a derivatised FA species. The exact mass (±0.01 Da) was then used to identify the number of carbons and double bonds present within the fatty acyl chain. The full chromatogram of each sample is displayed on the left, and a magnification of the region pertaining to ULCFA retention times is displayed on the right. Considering the longest FA species present within the standard mixture of 37 FAs contains 24 carbons, an absence of chromatographic features, such as FA 32:1 and 34:1, in the ULCFA region of the standard and the presence of longer retention times features with *m/z* values consistent with ULCFAs in the cell extracts provides further evidence that these ULCFAs are both present in the cell lines and are arising from biological means.

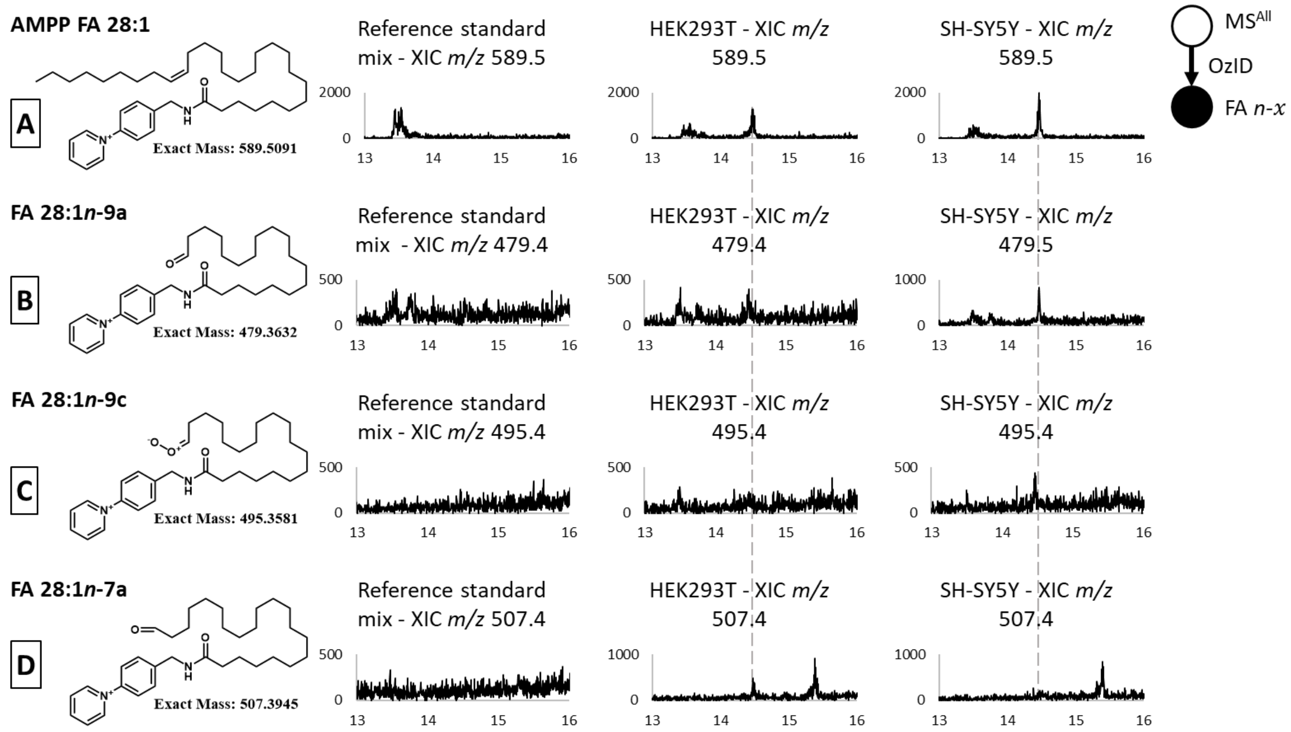


**Supplementary Figure 8.** Deep structural investigation and validation of the 28:1 fatty acyl using LC-MS-OzID. Chromatographically separated AMPP-FAs were exposed to ozone gas during their transmission through the ion-mobility cell of a modified Waters Synapt G2-Si mass spectrometer. This creates characteristic fragmentation of carbon-carbon double bonds and allows for double bonds position(s) to be determined through a neutral loss look-up table. Analogous to CID/OzID, OzID of olefins generates an aldehyde (row B) and Criegee (row C) product ion pair, which are used to validate the double bond positional assignment(s). Thus, temporal alignment between AMPP-FA precursor ions (row A) and OzID generated product ions (rows B-D) can signify the position of double bonds within the fatty acid. The absence of such product ion chromatographic features (indicated as chemical structures) from the reference standard mix provides further credibility towards the indicated OzID product ions stemming from the precursor ion. Rows B-D display that the FA 28:1*n*-9 and FA 28:1*n*-7 are present within the HEK293T and SH-SY5Y cell lines. (All ordinate axes are in counts per second, and abscissa axes are displayed in minutes).

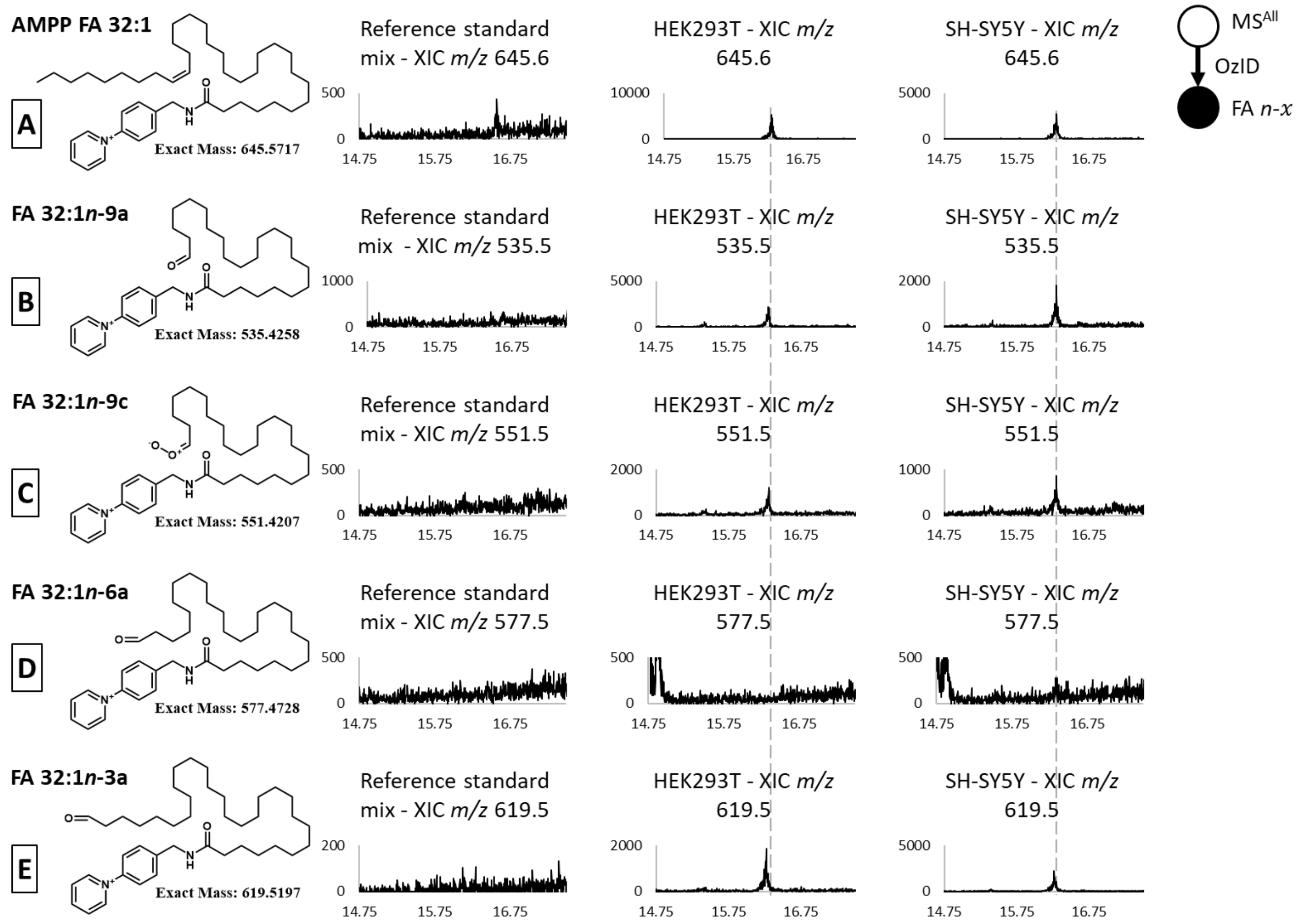


**Supplementary Figure 9.** Deep structural investigation and validation of the 32:1 fatty acyl using LC-MS-OzID. Chromatographically separated AMPP-FAs were exposed to ozone gas during their transmission through the ion-mobility cell of a modified Waters Synapt G2-Si mass spectrometer. This creates characteristic fragmentation of carbon-carbon double bonds and allows for double bonds position(s) to be determined through a neutral loss look-up table. Analogous to CID/OzID, OzID of olefins generates an aldehyde (row B) and Criegee (row C) product ion pair, which are used to validate the double bond positional assignment(s). Thus, temporal alignment between AMPP-FA precursor ions (row A) and OzID generated product ions (rows B-D) can signify the position of double bonds within the fatty acid. The absence of such product ion chromatographic features (indicated as chemical structures) from the reference standard mix provides further credibility towards the indicated OzID product ions stemming from the precursor ion. Rows B-E display that the FA 34:1*n*-9 and FA 34:1*n*-3 are present within the HEK293T and SH-SY5Y cell lines and FA 34:1*n*-6 is only present within SH-SY5Y. (All ordinate axes are in counts per second, and abscissa axes are displayed in minutes).

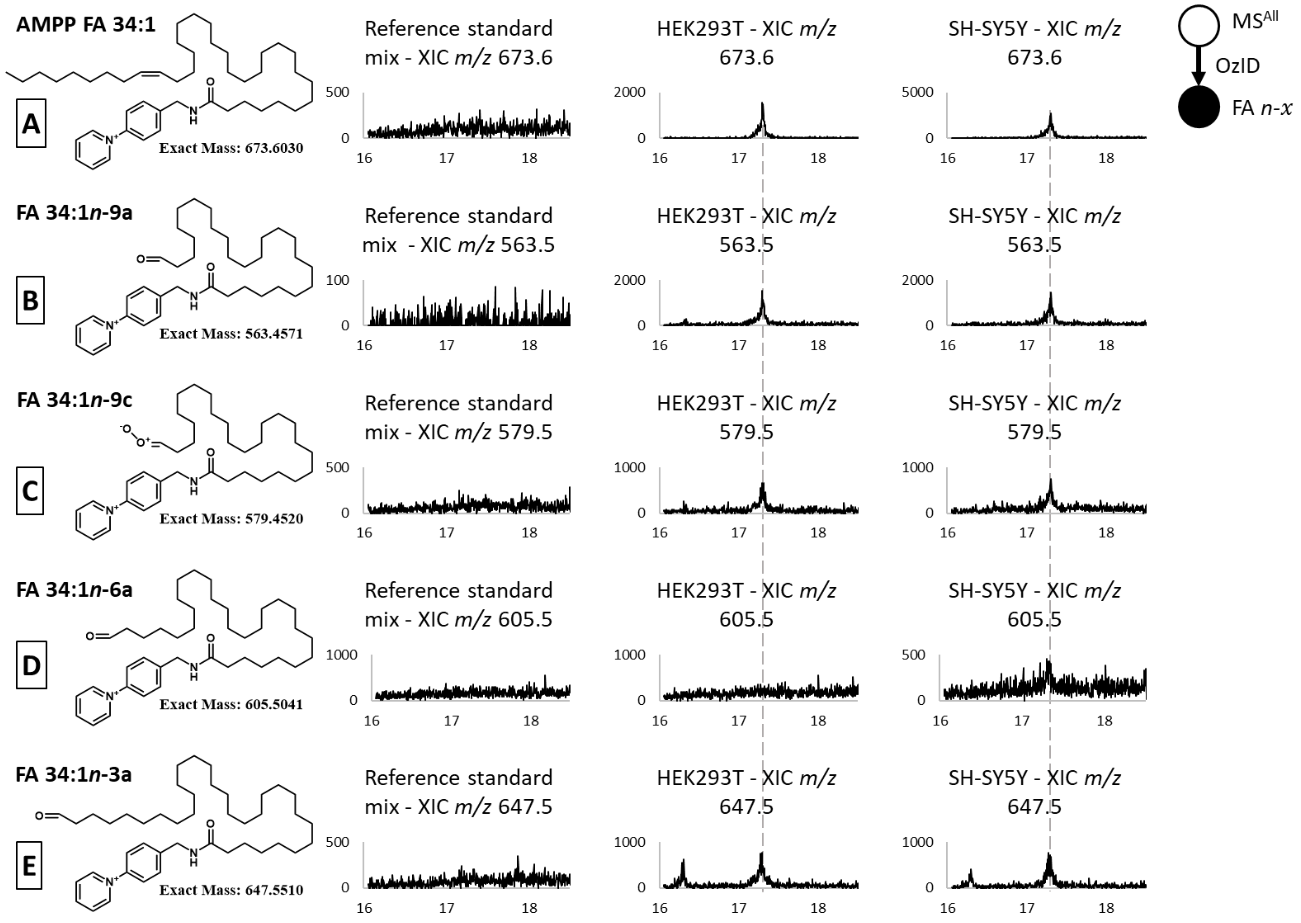


**Supplementary Figure 10.** Deep structural investigation and validation of the 34:1 fatty acid using LC-MS-OzID. Chromatographically separated AMPP-FAs were exposed to ozone gas during their transmission through the ion-mobility cell of a modified Waters Synapt G2-Si mass spectrometer. This creates characteristic fragmentation of carbon-carbon double bonds and allows for double bonds position(s) to be determined through a neutral loss look-up table. Analogous to CID/OzID, OzID of olefins generates an aldehyde (row B) and Criegee (row C) product ion pair, which are used to validate the double bond positional assignment(s). Thus, temporal alignment between AMPP-FA precursor ions (row A) and OzID generated product ions (rows B-E) can signify the position of double bonds within the fatty acid. The absence of such product ion chromatographic features (indicated as chemical structures) from the reference standard mix provides further credibility towards the indicated OzID product ions stemming from the precursor ion. Rows B-E display that the FA 34:1*n*-9 and FA 34:1*n*-3 are present within the HEK293T and SH-SY5Y cell lines and FA 34:1*n*-6 is only present within SH-SY5Y. (All ordinate axes are in counts per second, and abscissa axes are displayed in minutes).

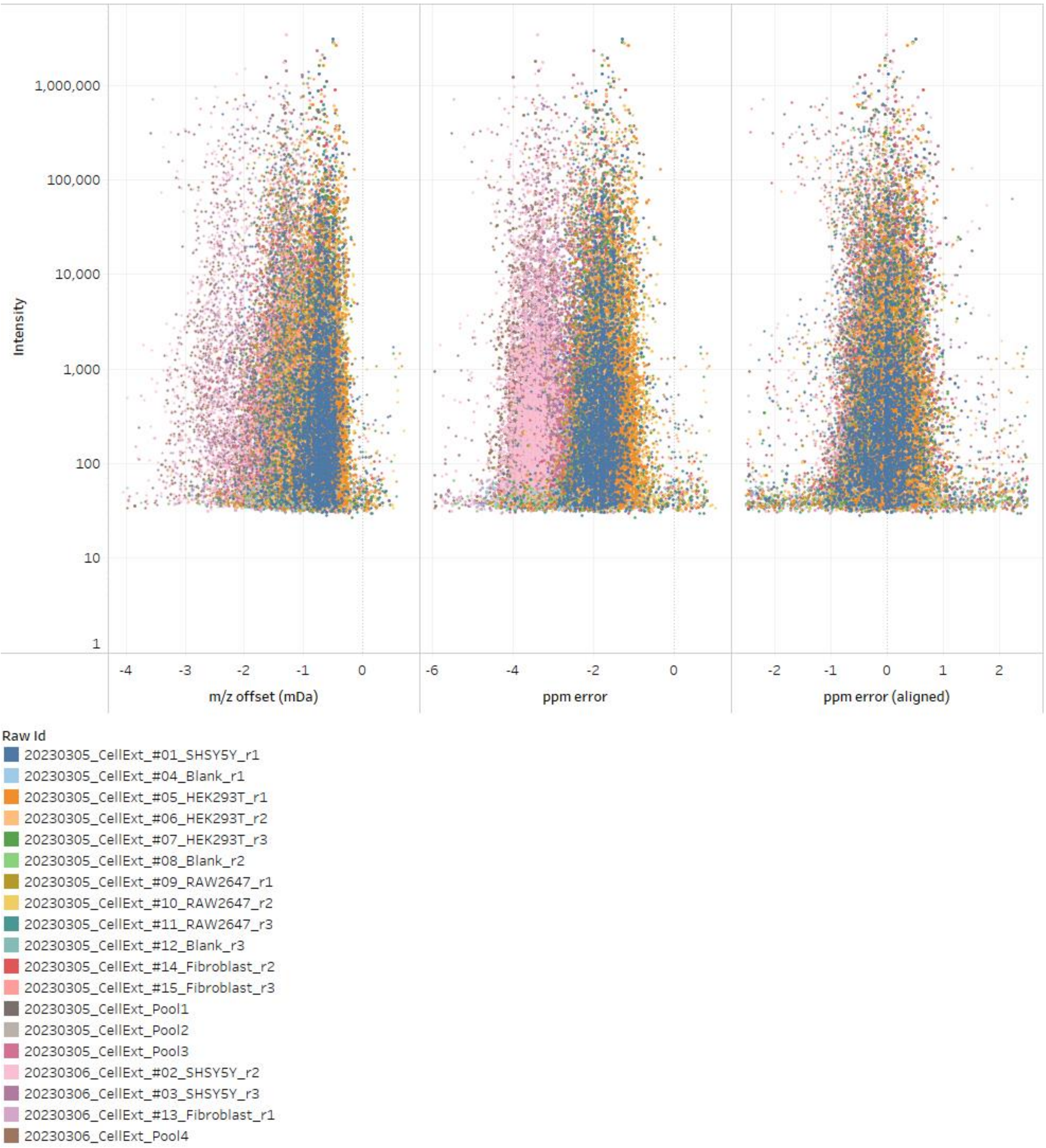


**Supplementary Figure 11.** Mass accuracy and influence of *m/z* realigment on CID/OzID fragment ion assignments. (left) *m/z* offset prior to realignment, (middle) ppm mass error prior to realignment, (right), ppm error after realingment. Realignment process is outlined in the methods. Each dot represents a single fragment ion assignment and each colour a different sample replicate data file.
